# Shared foraging grounds in a solitary rodent: indication for cooperation by kin selection and mutualism?

**DOI:** 10.1101/2024.07.07.602396

**Authors:** Lindelani Makuya, Neville Pillay, Carsten Schradin

## Abstract

1. Kinship is important for understanding the evolution of social behaviour in group living species. However, even solitary living individuals differentiate between kin and non-kin neighbours, which could lead to some form of cooperation, defined as both partners benefitting from each other. A simple form of cooperation is mutualism, where both partners benefit simultaneously.
2. Here we tested whether there is mutual tolerance by sharing foraging grounds between kin in a solitary species. This would indicate the possibility of kin selection and mutual cooperation.
3. We used mini-GPS data loggers to investigate range overlap in the solitary bush Karoo rat (*Otomys unisulcatus*) between kin- and non-kin neighbours. Next, we quantified the extent to which individuals shared foraging grounds containing food plants within their overlapping ranges. Lastly, using step selection functions applied to GPS fixes collected every five minutes, we analysed how individuals moved relative to each other.
4. Kin-neighbours had larger home range overlap than non-kin neighbours (70.4% vs 29.6%) and shared more of their foraging grounds (63% vs 37%).
5. Temporal analysis of spatial data found no indication that neighbours avoided each other, independent of kinship. Instead, activity was synchronised.
6. In sum, we found mutual tolerance between neighbours with regards to sharing foraging grounds, and kin shared nearly double as much of their foraging grounds than non-kin.
7. These data can be interpreted as a simple way of mutual cooperation between kin in a solitary species, where both members benefit from sharing a considerable part of their foraging grounds.

## Introduction

Solitary living species are not asocial, but often have non-random social interactions between neighbours which might even share resources (Makuya & Schradin 2024b). Solitary living refers to the social organisation (Kappeler & Schaik 2002) of a species where adult individuals live alone (Makuya & Schradin 2024a). Historically, it has been assumed that solitary living is caused by intra-specific aggression, functioning to monopolise resources (e.g. European hamster; *Cricetus cricetus*) (Eibl-Eibesfeldt 1953). However, a growing number of studies show that there are also amicable interactions between neighbours in solitary species, for example in mustelids (Twining *et al*. 2024), the African leopard (*Panthera pardus*) (Roex *et al*. 2022) and in estuarine crocodiles (*Crocodylus porosus*) (Baker *et al*. 2021). In the solitary puma (*Puma concolor*), individuals meeting at carrion respond in a non-random and consistent way, with some individuals always chasing each other aggressively, while other individuals commonly feed together (Elbroch *et al*. 2016; Elbroch & Quigley 2016). Whether such amicable sharing of food sources also occurs in solitary herbivores has not been studied in detail, though studies on kin selection and overlap in the kangaroo rats (*Dipomys ingens*) and Columbian ground squirrels (*Urocitellus columbianus*) suggest that sharing foraging grounds is possible (Meshriy, Randall & Parra 2011; Dobson *et al*. 2012).

The distribution of resources, such as food, influences the spatial distribution of animals (Johnson *et al*. 2002). The resource dispersion hypothesis states that if rare resources rare and evenly distributed, animals will be less likely to aggregate because there is no benefit for resource defence (Macdonald 1983; Johnson *et al*. 2002; Macdonald & Johnson 2015). In contrast, rich patchily distributed resources favour the formation of groups that defend these resources. Food distribution has been used to explain spatial structure of solitary species. For example, small, patchy food patches resulted in within group scramble competition and solitary living in the lemur *Microcebus berthae* (Dammhahn & Kappeler 2009). In contrast, the closely related lemur *M. murinus* relied on large patchily distributed food resources, thereby allowing females to overlap their home ranges and to form sleeping groups (Dammhahn & Kappeler 2009). While large patchily distributed resources drive group living in rodents (Ebensperger 2001), small patchily distributed food might drive solitary living to monopolize resources.

The distribution of resources influences social interactions between individuals (Halliwell *et al*. 2016). To understand the influence of resource distribution on social interactions, one must consider how individuals overlap spatially and temporally (Tania *et al*. 2012; Rodrigues 2018). Using step-selection functions on data obtained from GPS tracking, the movement of an individual can be related to the dynamic occurrence distribution of another individual, proving an estimation of fine-scale interactions (Schlägel *et al*. 2019; Ward *et al*. 2021). With this method one can determine whether individuals are attracted to conspecifics, avoid them or tolerate them. Fine-scale interactions have been quantified for bank voles (*Myodes glareolus*), where the males were always strongly attracted to females, while females were attracted to the males only when in oestrous but avoided them otherwise, maybe to avoid male infanticide (Schlägel *et al*. 2019). Similarly, the movements of male elongated tortoises (*Indotestudo elongata*) were driven by the presence of females (Ward *et al*. 2021). Both studies considered mates as a driver of the attraction of between individuals (Schlägel *et al*. 2019; Ward *et al*. 2021). Such fine-scale interactions between individuals could also provide insights into behavioural interactions underlying cooperation.

Cooperation between conspecifics has mainly been studied in group living species (Clutton-Brock 2016). The most basic definition of cooperation is that it consists of an interaction between two individuals that benefits both (Bergmüller *et al*. 2007a). Cooperation can evolve easily when both individuals benefit at the same time, referred to as by-product mutualism (Bergmüller *et al*. 2007a). An example can be seen in group living animals who gain safety from predators by forming large groups (Taborsky, Cant & Komdeur 2021). These animals cooperate by joint predator surveillance, whereby individuals take turns in watching for predators while the others forage, for example sentinels in meerkats (Clutton-Brock 2016). In such cases, where one individual first has costs before it later benefits, cooperation is either based on mutual reciprocity (Trivers 1971) or on kin selection leading to indirect fitness benefits (Hamilton 1971). Simple forms of cooperation might occur in solitary species that have a kin-based spatial structure with overlap in home ranges and frequent non-random interactions (Meshriy, Randall & Parra 2011; Makuya, Pillay & Schradin in press). Mutual tolerance is a requirement for cooperation and might even occur in solitary species, especially between close kin, adding indirect fitness benefits.

Most solitary mammals are either nocturnal, small and live in dense habitats, or very large predators roaming over large areas. Thus, field-based studies of species on both extremes are challenging. In contrast, the solitary and folivorous bush Karoo rat (*Otomys unisulcatus*) can be easily observed in its natural habitat (Schradin 2005; Makuya, Pillay & Schradin in press). Its endemic to semi-arid regions of South Africa and builds stick-lodges inside shrubs. It is a central-place forager, foraging within a short distance (less than 25m) from its lodge to patches of leafy succulent plants and ephemeral herbs, called foraging grounds, and then returning to the lodge (Schradin 2005). Runways lead from the lodge to the foraging grounds. Female bush Karoo rats form kin clusters, living closer to kin neighbours than to non-kin neighbours (Makuya, Pillay & Schradin in press). This makes it likely that foraging grounds of close kin overlap. Social interactions between close kin are characterized by tolerance while interactions between non-kin are more often aggressive, indicating a kin based amicable social structure (Makuya *et al*. 2024). The bush Karoo rat is therefore a good model to test the extent to which a solitary species shares patches of food and whether there is indication of kin selection and mutualism.

Here we asked whether the kin based spatial structure (Makuya, Pillay & Schradin in press) in combination with the amicable social structure (Makuya *et al*. 2024) might enable some simple form of cooperation (by-product mutualism) in the solitary bush Karoo rat. Specifically, we predicted (i) that foraging grounds will overlap more between kin-than non-kin neighbours, (ii) that neighbouring bush Karoo rats show positive interaction at foraging grounds and a (iii) higher temporal overlap of kin-compared to non-kin neighbours, indicating that kin use the same food resource at the same time while non-kin avoid each other.

## Methods

### Study site

Our study was conducted on a 4.5-hecare field site next to the Succulent Karoo Research Station located in the Goegap Nature Reserve, Northern Cape, South Africa (29°41’41.17” S 17°53’34.91” E). The area is a biodiversity hotspot, characterized by Succulent Karoo vegetation (Cowling, Esler & Rundel 1999; Myers *et al*. 2000). The climate is arid with minimum temperatures of -3°C in winter and maximum temperatures up to 43°C in summer (Schradin 2005). Mean annual precipitation at the field site is 160mm per annum. Seasons are characterised as the hot dry non-breeding season (November to April) and the cool moist breeding season (May to October).

### Trapping

Our field site was divided into six areas. Two areas were trapped simultaneously for three days by two people, before moving on to the next two areas. We used a combination of foldable Sherman traps and locally produced heavy traps of the Sherman style. Traps were placed at lodges that showed signs of being occupied (presence of fresh plant material, fresh faeces and active runways) by bush Karoo rats. All trapped rats were processed to obtain information on their weight (to the nearest gram) and their reproductive status (vagina perforated or closed; males scrotal or not). We conducted experiments using females as male bush Karoo rats disperse in autumn. We determined the kinship of the rats from trapping data: juveniles and pups were assumed to be the offspring of the adult female occupying the lodge where they were trapped; pups / juveniles were always trapped at the same lodge and during the breeding season only one adult female was ever trapped at a lodge at a time (Makuya, Pillay & Schradin in press). Upon first capture, rats were marked with metal band ear tags that had a unique identification number (National Band and Tag Co., Newport, KY, U.S.A.) (Schoepf & Schradin 2012). Rats were additionally marked with non-toxic hair dye to aid in individual identification during focal animal observations (Inecto Rapido, Pinetown, South Africa).

### Behavioural observations in the field

Focal observations were conducted five times a week at lodges with evidence of being occupied by bush Karoo rats to determine which individuals occupied the lodge (current study) and collect behavioural data for other studies (Makuya *et al*. 2024). The morning observations started five minutes before the sun started shining on the lodges and lasted for 30 minutes; afternoon observations of 30 minutes started 25 minutes before the sun stopped shining on the lodges. Three to five lodges were observed on the same day by different observers. Each lodge was observed for at least two days a month.

### Measuring daily ranges

Neighbouring adult female bush Karoo rats were simultaneously fitted with mini-GPS dataloggers (Gipsy 6 model from Technosmart, Italy https://www.technosmart.eu/gipsy-remote-copy/), from August 2021 to April 2023. We fitted 227 Gipsy collars on 43 days on a total of 125 females (on average 4.6 ± 1.5 females were fitted simultaneously, range 2 to 7 females). The mini-GPS loggers weighed 4.5g including the protective coating, which is less than 5 % of the rats’ body weight. These loggers were programmed to collect daily home range data according to methods validated by Makuya and Schradin (2023) and used by Makuya, Pillay and Schradin (in press). The mini-GPSs were fitted like a rucksack with a chest and a waist belt consisting of a wire running through a small rubber tube to avoid any skin irritation, fastened with a fishing crimp. The loggers collected the location (longitude, latitude, and altitude) of an individual and the date and time. We programmed the mini-GPS loggers to be off for the first two days (habituation) and start data collection on day 3 every 5 minutes; the battery life did not allow us to collect data for more than one day. We then re-trapped animals to remove the collars and to extract the data. Our method allowed us to assess whether movements of neighbouring rats are synchronised and to generate accurate estimate of distance travelled. A pilot study by Makuya showed that this method leads to similar home range sizes compared to using traditional method of collecting data every two hours for five days. The data were cleaned by first removing waypoints that had too high accuracy (> 50m) and outliers that were beyond 50m away from the centre.

### Measuring home range quality

Bush Karoo rats have runways leading from their lodges to foraging areas. We measured food availability along runways used by all female bush Karoo rats from which we had determined home ranges from the mini-GPS data. We ran a measuring tape along every runway within the ranges (Figure 1). Every 20cm, we recorded plants growing at 25cm and 50cm at right angles from the runways, both to the left and to the right. We recorded whether these plants were dead or alive. We further determined whether the plants were food plants based on data from nest observations. When there was no plant, a zero was recorded. For each runway, we counted the total number of food plants and associated the number of food plants to the runways that were then shared between the rats as determined from the GPS data.

**Figure 1:**
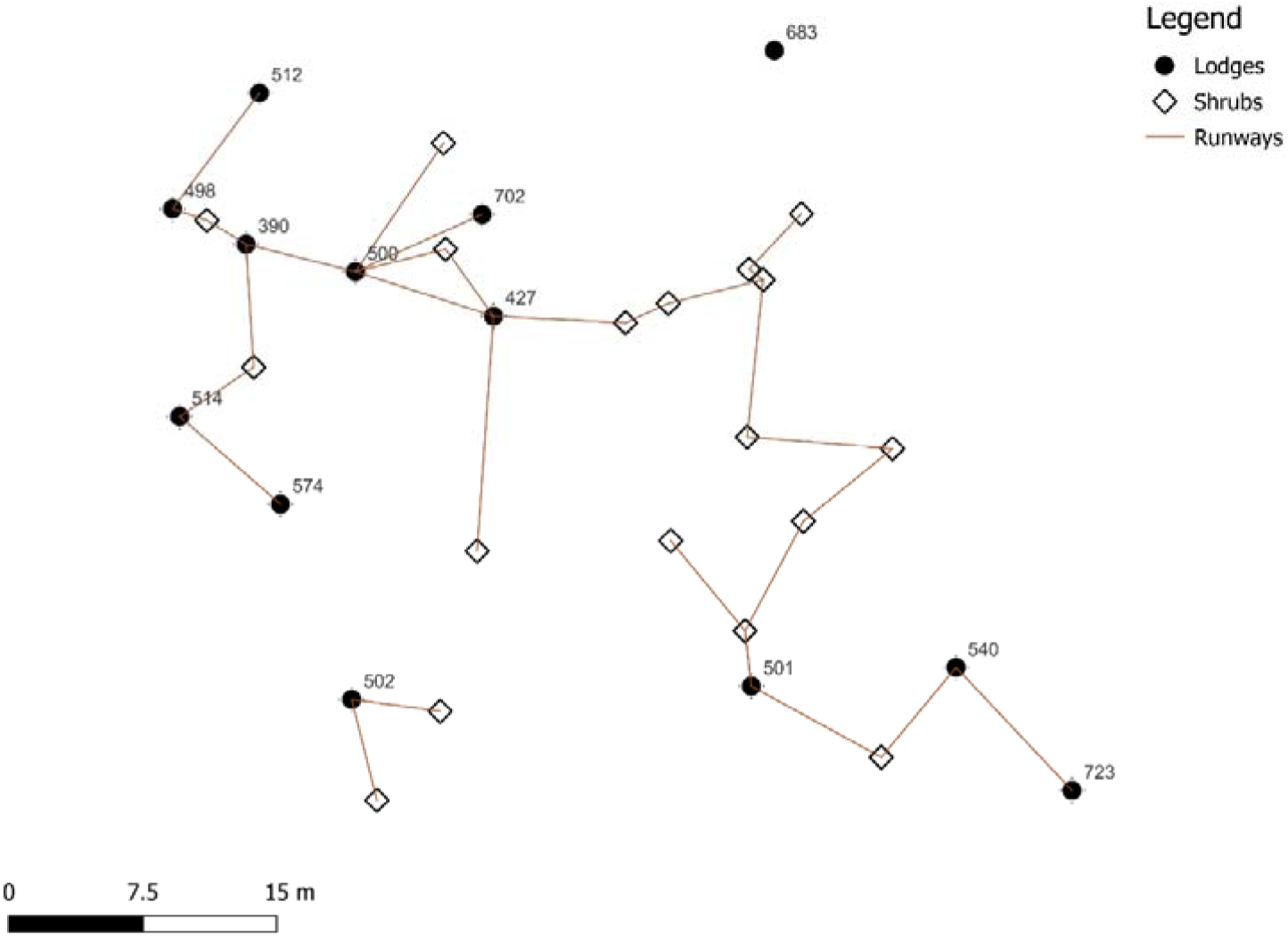
Example of lodges with runways. The hollow diamonds are the shrubs without lodges, black circles are lodges and brown lines are the runways between the lodges and shrubs.

### Statistical analysis

To analyse the daily range sizes and overlap, we used the autocorrelated Kernel density estimation (AKDE) implemented in the R packages *ctmm* (Calabrese, Fleming & Gurarie 2016) and *ctmmweb* (Dong *et al*. 2018). Overlap was calculated for every pair of rats (*N* = 192) on the daily range data from the same day. For the pairs of individuals that overlapped (*N* = 148), we estimated the interactions between individuals based on step-selection functions (SSF) in a workflow code supplied by Schlägel *et al*. (2019). This method allowed us to investigate whether the movement of one rat was correlated with the movement of its neighbour with which it overlapped. From this, we obtained a coefficient of interaction and p-value indicating whether there was attraction (positive value and *P* < 0.05; the rat chose locations where its neighbour also occurred), avoidance (negative value and *P* < 0.05: the rat avoided locations where its neighbour occupies) and neutral (close to zero coefficient: the rats’ choice of locations was independent of its neighbour’s occurrence). We then ran a general linear model (GLM) to analyse the factors that influence the coefficient of interaction. For this, we fitted the model (equation 1 below) using the fixed effects estimated overlap, relationship of the neighbour (kin vs non-kin) and season (breeding vs non-breeding) to determine whether these also had an impact on the response.

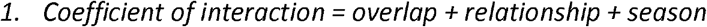

We determined the amount of food shared for all pairs of bush Karoo rats that overlapped in their ranges. For each pair, we determined which runways they shared and then added up all food plants counted along these runways. To analyse the factors that influenced the total shared food, we fitted a generalised linear mixed model (GLMM) with a Poisson distribution and a square root link function, run in the *lme4 R* package using the g*lmer* function (Bates *et al*. 2015), with the relationship (kin vs non-kin), season (breeding vs non-breeding) and the percentage of overlap fit as fixed effects. We also fitted the ID of both bush Karoo rats as random effects (equation 2 below).

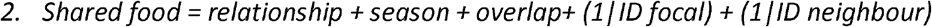

## Results

The home ranges of bush Karoo rats were larger in the moist breeding season (649.5 ± 424.3 m^2^, N = 82) than in the dry non-breeding season (447.6 ± 254.5 m^2^, N = 67; estimate= -405.4, SE = 446.5, p = 0.37). Their home ranges overlapped more with kin neighbours than non-kin neighbours (dry season: 49.8 ± 6.2 % vs 26.9 ± 4.7 % and wet season: 43.8 ± 5.3 % vs 21.6 ± 4.2 %) (relationship; estimate = -22.5, SE 5.3, *P* < 0.001; season; estimate = -5.7, SE = 5.3, *P* = 0.28).

### Overlap of foraging grounds

As predicted, we found higher overlap of foraging grounds between kin neighbours than non-kin neighbours (63% vs 37%, *P* < 0.001), especially during the dry, non-breeding season (dry season 66%, 38.5 ± 7.6 food plants for kin vs 33%, 26.04 ± 7.5 for non-kin; wet season 61%, 30.6 ± 6.9 vs 39%, 22.3 ± 6.1; *P* = 0.001, Table 1, Figure 2). The amount of shared food increased with increasing range overlap (*P* <0.001; Figure 3).

**Table 1:** Summary of Poisson GLMM to model the number of food plants shared by neighbouring bush Karoo rats.

| Model parameter | Estimate | SE | *P* |
| --- | --- | --- | --- |
| Intercept | -0.29 | 0.63 | 0.64 |
| Overlap | 0.069 | 0.004 | **<0.001** |
| Relationship (non-kin) | 1.85 | 0.33 | **<0.001** |
| Season (breeding) | 0.55 | 0.17 | **0.001** |

**Figure 2:**
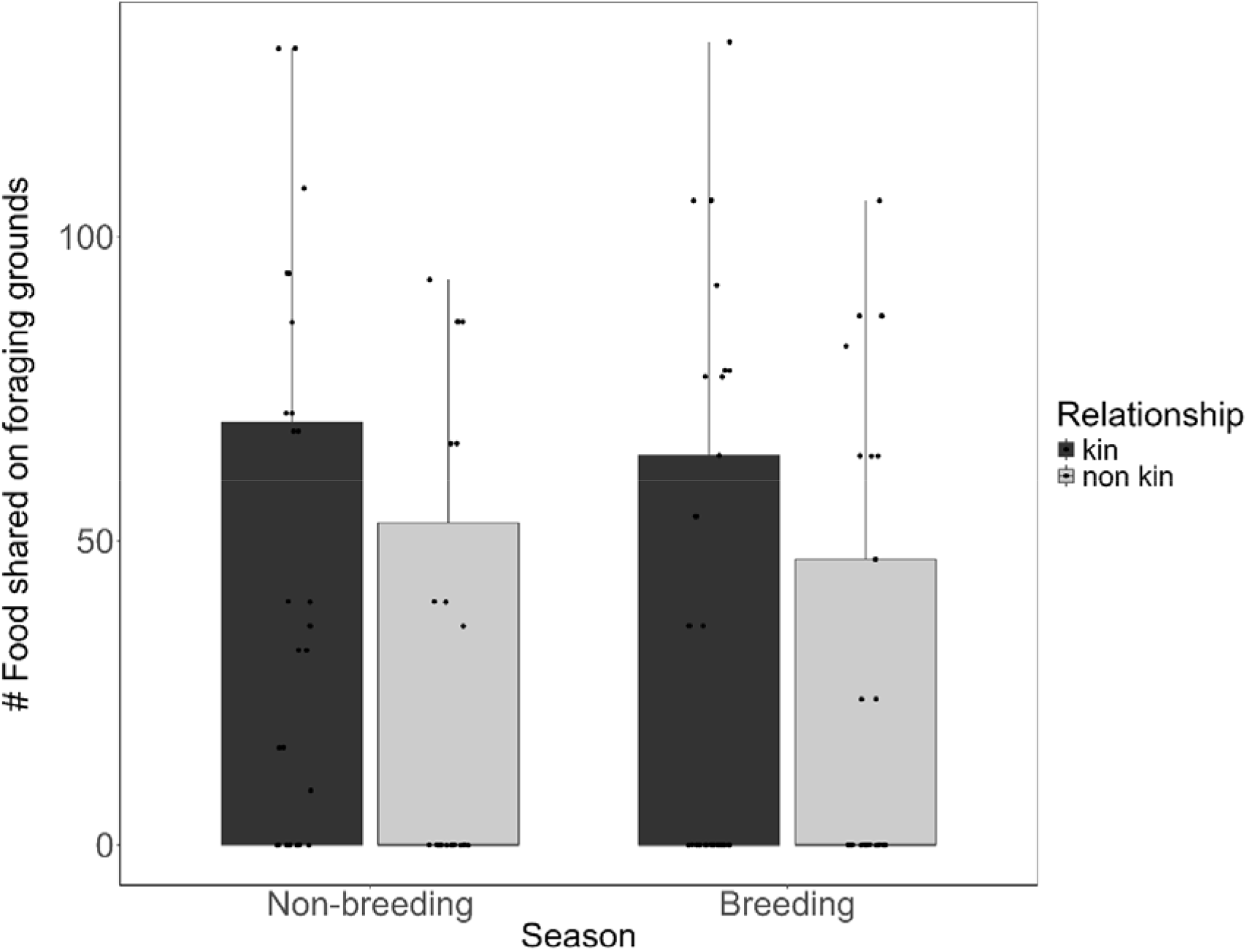
Seasonal comparison of total food shared between bush Karoo rats at foraging grounds by kinship. Boxplots show 1st and 3rd quartiles, the whiskers represent the minimum and maximum of the outlier data and points represent individual values.

**Figure 3:**
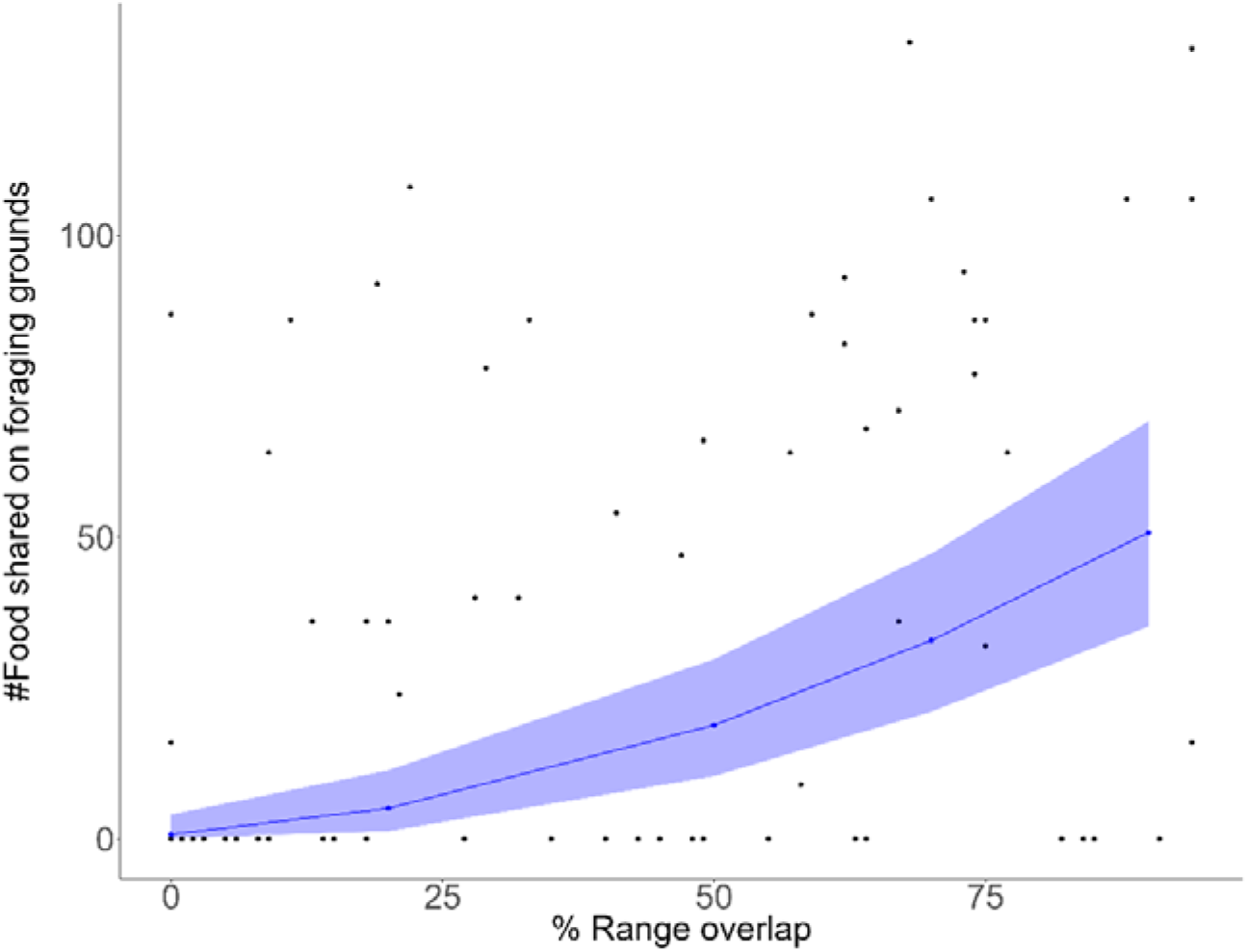
Mean fitted number of food plants within foraging grounds (solid line) and 95% confidence intervals (shaded area) against range overlap. Black dots are observed data.

### Temporal overlap (estimated step-selection coefficient of interaction behaviour)

The ranges of 148 pairs of bush Karoo rats overlapped between 2022 and 2023. From 90 of these pairs, we obtained useable temporal overlap estimates; for the remaining 58 pairs, the models either did not converge, or the model failed to calculate the coefficient of interaction because individuals only marginally overlapped. A total of 82 pairs were attracted to each other (coefficients greater than one), with 64 of these pairs being significantly attracted to each other (*P* < 0.05). Only three pairs avoided each other (negative coefficients) but not significantly (*P* > 0.05). Five pairs showed neutrality which was significant for only one pair (*P* < 0.05). Removing four outliers that had more than double the mean value for the coefficient of interaction, we therefore ran the GLM model for 86 pairs.

The coefficient of interaction was significantly and negatively influenced by the degree of range overlap (GLM; *t* = -5.60, *P* <0.001; Table 2 and Figure 4), i.e., rats which had a low overlap were more attracted to the same foraging grounds (higher coefficient) while rats which overlapped more were less or not attracted (neutral or negative attraction, coefficient close to zero). Unlike in our prediction, neither season nor relationship had a significant effect on the coefficient of interaction (GLM; *t* = 0.17, *P* = 0.869 and *t* = -0.33, *P* = 0.741 respectively).

**Table 2:** Summary of Gaussian GLM to model factors influencing the estimated step-selection coefficient (relative selection strength) of interaction behaviour in bush Karoo rats.

| Model parameter | Estimate | SE | *P* |
| --- | --- | --- | --- |
| Intercept | 8.7442 | 1.1378 | **<0.001** |
| Overlap | -8.3224 | 1.4862 | **<0.001** |
| Relationship (non-kin) | -0.2914 | 0.8779 | 0.741 |
| Season (breeding) | 0.1446 | 0.8757 | 0.869 |

**Figure 4:**
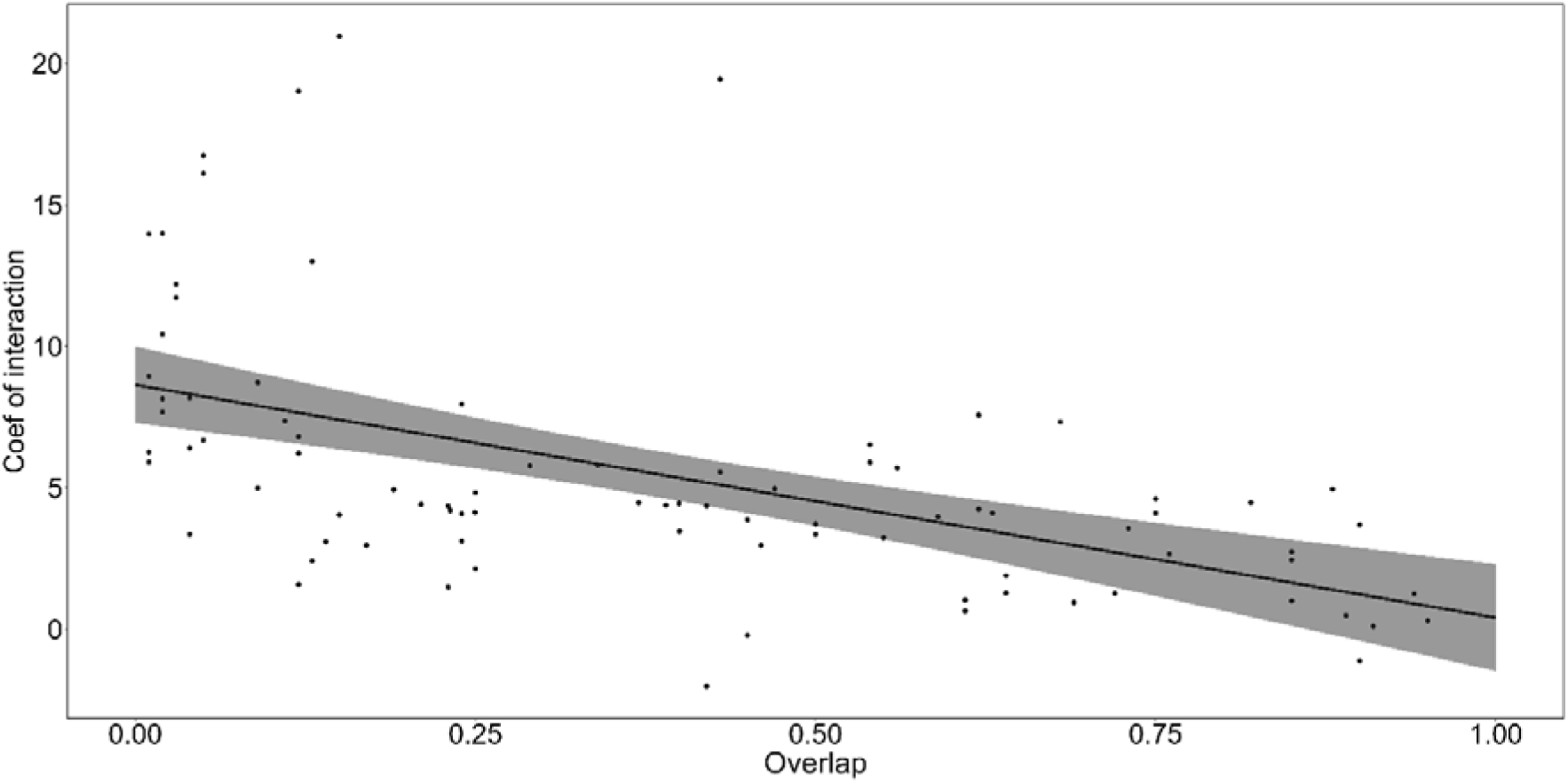
Mean fitted coefficient of interaction (solid line) and 95% confidence intervals (shaded area) against estimated range overlap. Black dots are observed data.

## Discussion

We tested whether closely related females of the solitary bush Karoo rat shared foraging grounds in a mutualistic way. Female bush Karoo rat shared more foraging grounds with their kin neighbours than their non-kin neighbours, and this especially in the food restricted dry season. They were attracted to each other at the foraging grounds, and we found no indication that they avoided each other. Therefore, they shared the restricted food resource mutualistically and especially between close kin, which can be interpreted as a simple form of cooperation.

The home ranges of female bush Karoo rats overlapped and so did their foraging grounds. We showed in a previous study that the overlap in home ranges was mostly driven by kinship (Makuya, Pillay & Schradin in press). Here we replicated this result: female kin shared nearly 50% of their home ranges, while overlap with non-kin was only half as much. Overlap in home ranges also lead to a significant overlap in foraging grounds. Female bush Karoo rat shared 61% of their food with their kin neighbours but only 39% with their non-kin neighbours in the moist season. In the food restricted dry season, when one might expect individuals to monopolize food and overlap thus to decrease, this difference was even stronger, and females shared 66% of their food with close kin but only 33% with non-kin.

Individuals can use the same resources without sharing them, for example by using them at different times of the day, or by avoiding each other and only visiting a resource when it its free. Therefore, we calculated interaction scores between dyads to assess whether they avoided each other. Interactions scores were positive for 90% of dyads independently of kinship, which could mean one of two things: either they were attracted to each other or they were attracted to the foraging grounds, an essential resource (Spiegel *et al*. 2016). Bush Karoo rats are central place foragers, which forage a short distance from their lodges to the foraging grounds. Previously we reported that they have as many kin as non-kin neighbours (Makuya, Pillay & Schradin in press) but behave differently to each of them. Female bush Karoo rats react with tolerance or are even amicable towards close kin, but are aggressive towards non-kin (Makuya *et al*. 2024). Together these results indicate that the positive interaction scores were rather due to attraction to the food within the foraging grounds than due to social attraction, in which case we would have expected positive association between kin and avoidance between non-kin. However, they did not avoid each other but visited the foraging grounds during the same time of the day, leading to positive interaction coefficients.

Bush Karoo rats shared more foraging grounds the more their home ranges overlapped, but at the same time their interactions scores declined, even though remaining positive in most cases. This indicates that they increased the distance between individuals when sharing larger areas. Thus, this effect might not indicate reduced competition but simply a statistical effect: when they share larger areas, by chance they are more often far away from each other.

Season had no influence on the interaction score, but on the number of shared food in the foraging grounds. If sharing of food is simply a consequence of overlapping areas with high food availability (food being an abundant resource that does not need to be defended), one would have expected them to share more foraging grounds in the food rich moist season and to monopolise food when it becomes rare by decreasing overlap. Surprisingly, we found the opposite: more overlapping foraging grounds in the food restricted dry season. This further indicates that tolerance between female bush Karoo rats did not decrease when food became scarce. Indeed, in another study, we observed that aggression between neighbouring females is higher in the food rich breeding season compared to the dry season, probably to avoid reproductive competition such as female infanticide (Makuya *et al*. 2024). This indicates that food is only a minor driver of competition and explains why foraging grounds can be shared in a mutualistic way.

By-product mutualism is a form of cooperation where both parties benefit at the same time (Bergmüller *et al*. 2007b). However, it is only regarded as a form of cooperation if it includes an interaction between the two parties (Bergmüller *et al*. 2007b), especially if the behaviour involved was under selection for this purpose (West, Griffin & Gardner 2007). Whether mutual sharing of foraging grounds in bush Karoo rats represents a simple form of cooperation is not completely evident: while both parties potentially benefit from having access to the foraging grounds, this is not characterised by interactions but rather by social tolerance allowing both individuals to exploit this resource. Importantly, tolerance is not a random response but represent an absence of aggressive interactions. This indicates that tolerance is an evolved trait that avoids aggressive interactions, and could thus also be regarded as a kind of neutral interaction. For example, in solitary carnivores, some individuals do regularly tolerate each other when foraging at carcasses, but aggressively chase away other individuals from the same food source (Elbroch & Quigley 2016; Roex *et al*. 2022; Twining *et al*. 2024). Some degree of tolerance is necessary for all group-living species, which do not automatically classify as cooperative because they share foraging grounds, but typically only when they directly interact to share food (Clutton-Brock 2002). Thus, in several solitary mammals, individual-specific tolerance for mutual sharing of food resources has evolved, often based on kinship, indicating early forms of by-product mutualism

## Conclusion

In conclusion, the spatial structure has an influence on the social interactions between individuals. Specifically, solitary bush Karoo rats overlapped more with their kin neighbours and therefore shared more foraging grounds, without avoiding each other. This adds to the growing number of studies that indicate that solitary species can have non-random social interactions with their neighbours. Attraction to the same foraging grounds leads to the sharing of food, especially between kin, and the mutual tolerance of each other together with the synchronised behaviour could be interpreted as an interaction that benefits both, or at least as the absence of an aggressive interaction to monopolise food, and as such as a simple form of cooperation. Such forms of cooperation could be the starting point for the evolution of more complex forms of cooperation including amicable interactions.

